# What Drives Symbiotic Calcium Signalling In Legumes? Insights And Challenges Of Imaging

**DOI:** 10.1101/559971

**Authors:** Teresa Vaz Martins, Valerie N. Livina

## Abstract

We review the contribution of bioimaging in building a coherent understanding of Ca^2+^ signalling during legume-bacteria symbiosis. Currently, two different calcium signals are believed to control key steps of the symbiosis: a Ca^2+^ gradient at the tip of the legume root hair is involved in the development of an infection thread, while nuclear Ca^2+^ oscillations, the hallmark signal of this symbiosis, controls the formation of the root nodule, where bacteria fix nitrogen. Additionally, different Ca^2+^ spiking signatures have been associated with specific infection stages. Bioimaging is intrinsically a cross-disciplinary area that requires integration of image recording, processing and analysis. We use experimental examples to critically evaluate previously established conclusions and draw attention to challenges caused by the varying nature of the signal-to-noise ratio in live imaging. We hypothesise that nuclear Ca^2+^ spiking is a wide-range signal involving the entire root hair, and that Ca^2+^ signature may be related to cytoplasmic streaming.

## 1. Introduction

Nitrogen is the most common limiting nutrient for plant growth, but a particular class of plants, legumes, can overcome this limitation with the help of bacteria [1,2]. Legumes secrete flavonoids, which are recognised by soil bacteria called rhizobia. In response, rhizobia release nodulation (Nod) factors (NF). This exchange of signals marks the beginning of a symbiotic relationship. Typically, the root hair tip swells and curls to entrap the bacteria (see Fig. 1), which then enter the root upon the formation of a plant-made tunnel-like structure called the infection thread. This allows the bacteria to cross several cell layers. The process culminates in the formation of a nodule, where bacteria will fix nitrogen for the plant in exchange for sugars. Throughout rhizobial infection and the organogenesis of the nodule, the generation and decoding of biochemical signals tells the plant what to do, and central to this process are Ca^2+^ signals.

**Figure 1.**
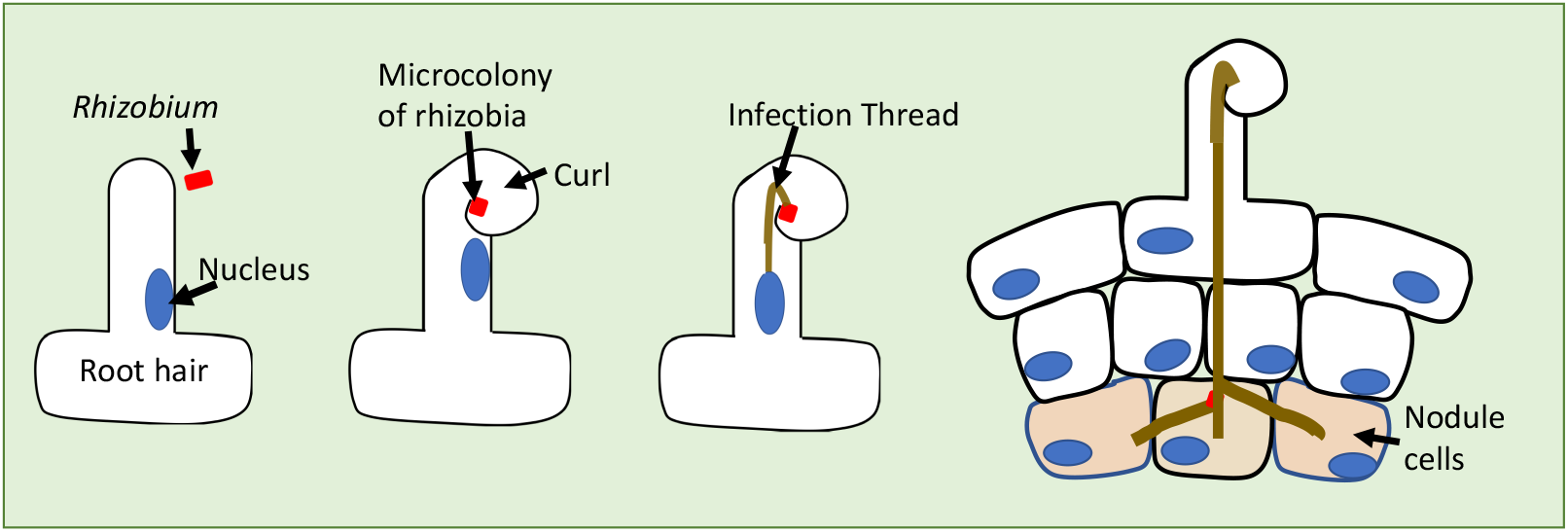
Diagram of the response of a *M. truncatula* legume root to nitrogen-fixing rhizobia. After exposure to rhizobial signals - Nod factors - the legume root hair curls to entrap the bacteria, which then penetrate into the root via a plant-made infection thread. Branches of the infection thread penetrate cortical cells, which proliferate leading to the formation of a root nodule to house the bacteria. Nodule formation takes place over the course of several days, while imaging is usually done only for a few hours after root hairs exposure to Nod factors. Ca^2+^ signals have been linked to the infection thread growth and nodule formation by analysis of different mutants. The blue oval denotes the nucleus, which moves to guide the formation of the infection thread [3,4], The nucleus is anchored by cytoplasmic strands which connect it to the cytoplasm (not represented, it presents different cytoarchitecture according to the root hair development stage).

Investigations into the role of symbiotic calcium signalling integrate knowledge from different disciplines, from genetics to electrophysiology, mathematical modelling and bioimaging. Traditionally, the detection of Ca^2+^ signals relied on electrophysiological methods, but microelectrodes can only measure calcium at one cellular point. With the development of fluorescent calcium sensors [5–9], live imaging has revealed diverse spatio-temporal Ca^2+^ dynamics. This has profoundly impacted the field, as it indicated where to look for the components of the Ca^2+^ signalling machinery. This inspired mathematical models [10,11], which attempted to explain the generation of Ca^2+^ signals.

The goal of this review is to revisit the contribution to current knowledge about Ca^2+^ signals during legume-bacteria symbiosis, from the point of view of bioimaging. Various reviews describe the field from different angles [1,2,12–15], Bioimaging is special in that it crosses different disciplines and is now being taken to a new level by crucial advances in imaging processing and quantification [16,17], It is timely to revisit the imaging evidence obtained so far and identify blind spots that demand a fresh look. Here we will focus only on the conclusions drawn from fluorescent microscopy.

Rhizobia commonly invade legumes via their root hairs, as shown in Fig. 1. In this case, the primary regions of interest are the root hair tip that curls to entrap bacteria, and the nucleus where transcription takes place. Bioimaging suggests that there are distinct Ca^2+^ signals predominantly associated with each region: a Ca^2+^ influx at the tip region, which is thought to be involved in the formation of an infection thread, and nuclear Ca^2+^ oscillations, which are required for infection thread growth and nodule organogenesis. Evidence from time-lapse imaging supports the concept of Ca^2+^ signatures [18–21], such that representative Ca^2+^ patterns can be associated with different mutants, infection stages, or outcomes.

It is not trivial to compare Ca^2+^ signals in different regions of the cell. Genetically encoded Ca^2+^ sensors (GECI) [22] are increasingly popular largely because, in comparison with dyes, they are easier to target in different cellular compartments. One challenge is that the detected fluorescence depends on the fluorophore concentration, which in turn depends on the cell thickness and cellular motions in and out of focus: an example is root hair cytoplasmic streaming, which changes patterns as the root hair grows, causing changes in the distribution of cytoplasmic sensors. To compare Ca^2+^ transients in different cellular regions, researchers measure ratios instead of absolute values [23]. This can be done by using ratiometric indicators that shift the absorption or emission wavelength upon calcium binding, such as Fura-2 or calcium cameleons [24,25], or by making pseudo-ratiometric measures when co-loading a non-ratiometric indicator such as Calcium Green or Oregon Green with a Ca^2+^-insensitive reference dextran, such as Texas Red.

In any image, the Ca^2+^ signal must be detected over the background of fluorescent noise, generated by the detector or arising from the random nature of photon release [26]. The overall ability to detect the signal can be determined by considering the ratio of the signal to noise levels of the image [27]. Whether ratiometric or not, Ca^2+^ indicators are only effective if there is sufficient signal-to-noise ratio (SNR). Although some characteristics of Ca^2+^ signals have been confirmed using several different sensors, imaging processing methods have not been validated by independent approaches, as reliable signal processing methods are still in their infancy in this field. This review will provide examples of how implicit assumptions in image processing and analysis can bias the images we see.

## 2. Two symbiotic calcium signals

In the next two subsections 2.1 and 2.2, we will review the Ca^2+^ signals responses currently associated with two processes: formation of an infection thread and nodule organogenesis (Fig. 1). Table 1 lists some papers that have informed the current view of two separate Ca^2+^ signals, and the Ca^2+^ indicators that have allowed a comparison. We distinguish two basic different signals. Ca^2+^ oscillations are predominantly associated with the nuclear region, and calcium gradients with the tip region. This is a simplified view, as there are areas of intersection: both in terms of response patterns, and in terms of location.

**Table 1.**
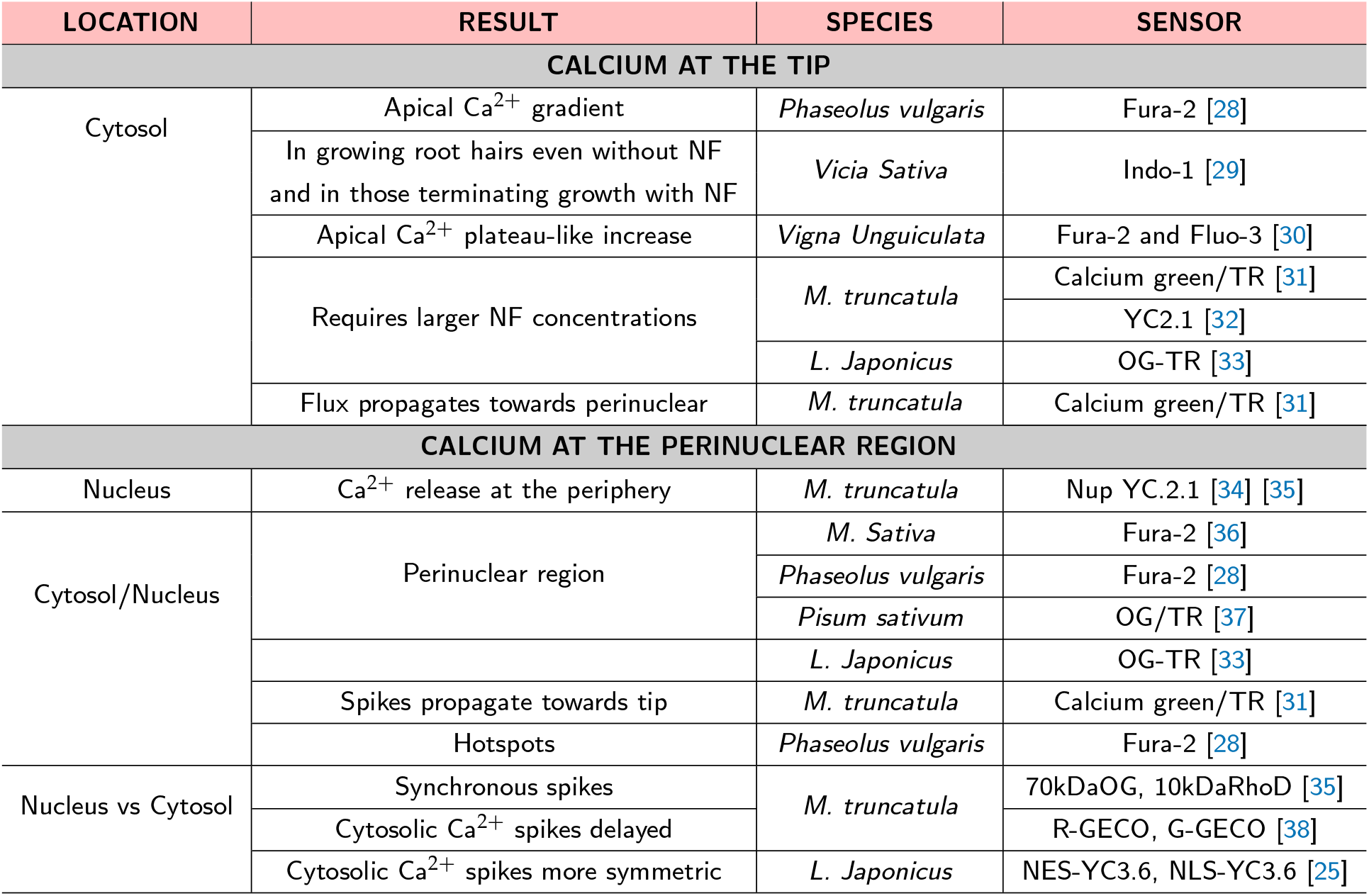
Two different Ca^2+^ signals induced by Nod factors (NF) were detected in several model legumes, with different Ca^2+^ sensors: dyes (Fura-2, OG, Indo-1, Fluo-3, RhoD) and genetically-encoded sensors (the yellow cameleons YC2.1 and YC3.6, and the GECOs R-GECO and G-GECO)

### 2.1. Ca^2+^ levels rise at the root hair tip

An increase in Ca^2+^ concentration at the root hair tips is a common signalling response to Nod factor [28–33,37,39] in various legumes, Table 1. Combining evidence from imaging with the design of mutants with different impairments, it is suggested that apical Ca^2+^ flux is related to the activation of the growth of the infection thread [33]. Imaging studies support this picture on several grounds. While an apical Ca^2+^ gradient is observed in developing root hairs even in the absence of Nod factor [29], the root hairs that are terminating growth respond to Nod factor not only by a Ca^2+^ flux, but also by recovering additional characteristics associated with tip-growing cells, namely a polar cytoplasmic organisation and reverse fountain streaming. Notably, it is at this stage of development that the root hair is more susceptible to rhizobial infection [40]. Moreover, a Ca^2+^ flux response requires increased Nod factor concentration [31] compared to the other symbiotic Ca^2+^ response (Ca^2+^ spiking, see Subsection 2.2). This is a suggestive piece of evidence because in nature larger Nod factor concentrations occur when bacteria become entrapped in a enclosed curled root hair.

In imaging studies, a calcium flux is typically the first Ca^2+^ signalling response, appearing within 1-5 minutes after Nod factor addition. [28,31,37]. But in nature, morphological events such as root hair curling and infection thread growth take place hours and not minutes after exposure to bacterial Nod factors. However, Ca^2+^ flux appears as an earlier response only when Nod factor is applied in large concentrations from the start. If the concentration is increased later, the response can be induced in root hair cells that are already spiking [31,37]. Therefore, imaging evidence is compatible with a scenario in nature where Ca^2+^ oscillations would occur earlier and a Ca^2+^ flux could happen later when Nod factor concentration is raised by the increasing number of rhizobia growing within the compartment generated by a curled root hair.

The Ca^2+^ flux is usually described as a biphasic response with a single rapid rise in cytoplasmic calcium concentration followed by a slower decline to baseline calcium levels. In the context of the symbiosis between legumes and bacteria, a different signal, Ca^2+^ oscillations, is conventionally associated with another root hair region, the perinuclear region, as we will see in Subsection 2.2.

### 2.2. Ca^2+^ spiking around and inside the nucleus

Whilst apical Ca^2+^ signals are associated with growth in several systems [41,42] and cytosolic Ca^2+^ signals participate in processes like plant pathogen defence, nuclear calcium oscillations are a distinctive feature of symbiotic calcium signalling [13,43], genetically correlated with nodulation [36,37,44]. Regular oscillations in Ca^2+^ concentration (Ca^2+^ spiking) have been observed in legume root-hair cells, approximately ten minutes after exposure to Nod factor. Ca^2+^ spikes are asymmetric, with a rapid rising phase followed by a gradual decline [36] (see the golden trace in Fig. 5, panel A). Although adjacent cells don’t spike at the same time [25,34], they mostly share typical calcium oscillations, with similar frequencies and spike shapes. As it is typical in plants, the kinetics of Ca^2+^ dynamics are much slower than in the animal field [45]. In the model legume *M. truncatula*, the typical Ca^2+^ oscillations periods vary between 50 and 100 seconds.

Perinuclear Ca^2+^ spiking is a common response to rhizobial signals in legumes. It was observed at pre-infection stages in legumes that form indeterminate nodules, such as *Medicago truncatula* [44,46,47], *Medicago sativa* (alfalfa) [36] or *Pisum sativum* (peas) [37], as well as in legumes like *Phaseolus vulgaris* (beans) [28] and *Lotus japonicus* [33,48] that form determinate nodules. It was also observed in *Sesbania rostrata* [49], a legume which can undergo intercellular lateral basal bacterial invasion in addition to the root hair invasion. Perinuclear Ca^2+^ spiking persists for some time during trans-cellular infection, as confirmed in *M. truncatula* [50].

In all cases the oscillations appear sharper near the nuclear area [36,37], tending to become undistinguishable with the distance from it: when calcium spikes are visible at the tip, the spike rising phase appears more gradual [25]. Possibly [25], this difference reflects Ca^2+^ release from sources at the nuclear region, and its dissipation as it propagates towards the root hair tip. An alternative hypothesis is that since the cytosol is disproportionally concentrated around the nucleus [37], farther from the nucleus the fluorescent cytosolic Ca^2+^ signal becomes masked by noise. Naturally, the independence of ratiometric techniques from the sensor distribution is only valid assuming that the concentration of the sensors is large enough to allow a good signal-to-noise ratio.

To directly look for the origin of Ca^2+^ signals, Shaw and Long [31] measured delays between Ca^2+^ signals in different cellular regions. Comparing Ca^2+^ signals imaged at one-second intervals, they show that Ca^2+^ spiking starts near the nucleus. A similar increasing delay of Ca^2+^ spikes with distance from the nucleus was reported by Kelner et al [38]. Moreover, perinuclear Ca^2+^ spiking can occur without a previous apical Ca^2+^ flux [31,33]. Therefore, to find the origin of perinuclear spiking, it seemed natural to look around its vicinity, starting from inside the nucleus itself [34].

Whilst several of the dyes previously used didn’t discriminate between nucleus and cytosol [43], Sieberer et al [34] used a genetically encoded yellow cameleon, NupYC2.1, specifically targeted to the nucleus. They reported sustained and regular intranuclear Ca^2+^ spiking in the nuclei of *Medicago truncatula* root hairs treated with NFs. In terms of frequency and asymmetric spike shape, the characteristics of intranuclear Ca^2+^ oscillations were similar to those [36] of perinuclear cytosolic Ca^2+^ signals. Spatially, they observed that the intranuclear Ca^2+^ concentration was unevenly distributed, and they suggested that each spike originated from the periphery of the nucleus. That could indicate a nuclear envelope Ca^2+^ store, and the idea is reinforced by similar imaging results with pseudo-ratiometric indicators [35], as well as by the localisation of the components of a calcium signalling machinery in the nuclear membranes [35,51,52]. However, Sieberer et al [34] note that the uneven distribution of intranuclear Ca^2+^ could result from the signal-free nucleolus, but also indicate the presence of possible intranuclear release sites. Regardless of its spatiotemporal signature, the main result in Sieberer et al [34] was to confirm the existence of intranuclear Ca^2+^ spiking.

Since nuclear and cytosolic signals have quite similar time dynamics [35], the nucleus is porous [53], and the nuclear envelope consists of a double membrane with some components of the Ca^2+^ signalling machinery on both the cytosolic- and the nuclear-facing side [35,52], cytosolic and nuclear Ca^2+^ signals might be coordinated. Whilst a previous study [35] could not discriminate between a cytosolic and nuclear Ca^2+^ origin, recently [38] an important step was made with the development of new genetically-encoded calcium sensors, GECOS, with a much improved dynamic range. Using a dual colour sensor able to simultaneously detect cytosolic and intranuclear Ca^2+^ signals, Kelner et al [38] report that the majority of perinuclear cytosolic Ca^2+^ spikes starts with a delay relative to nuclear Ca^2+^ spikes. However, it was not possible to clearly distinguish between nuclear and cytosolic Ca^2+^ spiking initiation sites in 30% percent of the cases, and in a very few cases (2%) intranuclear Ca^2+^ spikes appeared delayed relative to perinuclear Ca^2+^ spikes. In time-lapse live imaging, it can be very challenging to delineate boundaries of small regions. In the study [38], the authors chose an oval as the region-of-interest (ROI) to represent the nucleus, while the perinuclear region was represented by an annular-shaped ROI around the nucleus. Both ROIs appeared static, which can potentially lead to partial loss of signal if the nucleus or the root hair move slightly, especially for the case of the narrow annular-shaped perinuclear ROI. Perhaps the percentage of inconclusive results could be settled by the choice of larger/dynamic ROIs for the perinuclear region. In case the thickness and position of the confocal slice allows the observation of overlapping nuclear and cytosolic signals, the nuclear and perinuclear ROIs can even coincide, taking advantage of the ability of the sensor to discriminate between nuclear and cytosolic signals.

## 3. Calcium signatures

Bioimaging has shown that Ca^2+^ signals can adopt rich spatio-temporal patterns. In some plants and animals, calcium spatio-temporal patterns can encode information about the primary stimuli and the attribution of coding properties to calcium signals inspired the search for representative calcium traces. Among other characteristics, the so-called *calcium signatures* can have different frequencies or spike shapes, and can be typical of different root hair development stages [46], infection stages [50] or different bacterial invasion strategies [49]. Table 2 lists some of the biological phenomena in rhizobial-legume symbiosis that have been associated with different Ca^2+^ signatures.

**Table 2.**
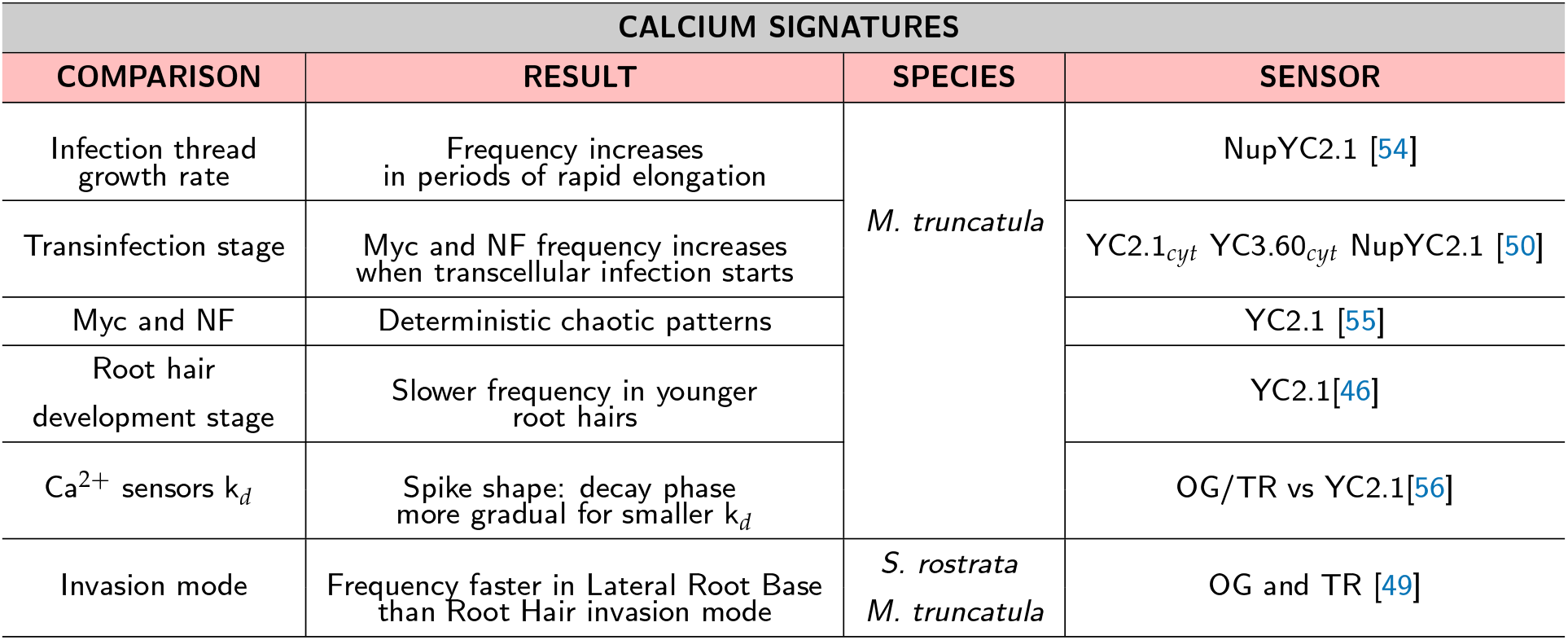
Calcium signatures: different Ca^2+^ oscillation frequencies or spike shapes characterise different phenomena.

| CALCIUM SIGNATURES |  |  |  |
| --- | --- | --- | --- |
| COMPARISON | RESULT | SPECIES | SENSOR |
| Infection thread growth rate | Frequency increases in periods of rapid elongation | *M. truncatula* | NupYC2.1 [54] |
| Transinfection stage | Myc and NF frequency increases when transcellular infection starts |  | YC2.1<sub>cyt</sub> YC3.60<sub>cyt</sub> NupYC2.1 [50] |
| Myc and NF | Deterministic chaotic patterns |  | YC2.1 [55] |
| Root hair development stage | Slower frequency in younger root hairs |  | YC2.1[46] |
| $\text{Ca}^{2+}$ sensors $k_d$ | Spike shape: decay phase more gradual for smaller $k_d$ | | OG/TR vs YC2.1[56] |
| Invasion mode | Frequency faster in Lateral Root Base than Root Hair invasion mode | *S. rostrata*<br>*M. truncatula* | OG and TR [49] |

### 3.1. Ca^2+^ frequency

In legume-microbe symbiosis, calcium signals are part of a biochemical signalling pathway, with several components shared with another symbiosis that most plants establish with mycorrhizal (Myc) fungi to meet their demands for phosphorus. An attractive hypothesis is that Ca^2+^ signals encode specific information about the primary stimuli, either fungal Myc factors or bacterial Nod factors. However, although one study [55] suggested that Myc and Nod factors induce different Ca^2+^ signatures with a flexibility provided by their deterministic chaotic nature, the authors noted that different experimental approaches may have played a part in the results. And while doubts remain about the specificity of Ca^2+^ signatures during pre-infection, at later symbiosis stages the patterns of Ca^2+^ oscillations are similar in both Myc and Nod symbioses, and related instead to the stage of infection [50]: in both symbioses, Ca^2+^ spiking switches from infrequent and irregular, to a pattern of high-frequency regular oscillations when transcellular infection begins [50].

In a study by Sieberer et al [50], the period of rapid spiking associated with apoplastic cell entry lasted for a total of 35-45 consecutive spikes, a similar number to the about 36 consecutive Ca^2+^ spikes predicted to be critical for regulating the ENOD11 gene activation involved in nodulation [46]. In fact, simple mathematical models [57] propose explanations for decoding mechanisms based, among other features, on the number of spikes. Wheareas Ca^2+^ spiking stops before transcellular infection has finished [50], it lasts for the entire duration of bacterial progression through the infection thread at the root hair [54], with faster frequencies when the rate of elongation is more rapid. If the growth of the infection thread stops, nuclear Ca^2+^ spiking is not observed [54].

Besides being related to infection stages in bacteria-legume symbiosis [50,54], the frequency of Ca^2+^ oscillations increases with the age of the root hair [46], and is also related to the bacterial mode of invasion. Bacterial infection via root hair invasion is common in legumes, but a few legumes can also nodulate at lateral root bases via a form of crack entry. Such is the case for the semi-aquatic legume *Sesbania rostrata*, capable of experiencing either mode of invasion. While oscillations associated with root hair invasion on growing root hair cells are comparable in *S. rostrata* and *M. truncatula* [49], the frequency of perinuclear calcium oscillations is faster and each spike more symmetric when *S. rostrata* is nodulated at lateral root bases.

Note that despite of the notion of characteristic oscillation frequencies, an oscillation frequency may vary with time. For instance, the onset of a more or less regular period of Ca^2+^ oscillations is often preceded by a period of initial rapid spiking, which can be stimulated again by repeated addition of Nod factor [56].

### 3.2. The shape of calcium spikes

A typical symbiotic calcium spike induced by Nod factor is asymmetric [36], with a rapid rising phase (1-4 seconds) followed by a more gradual decline that lasts more than 30 seconds. This is thought to correspond to the opening of calcium release channels, followed by the slower pumping of calcium back into Ca^2+^ stores.

As expected, the speed of the decay phase depends on the equilibrium dissociation constant (K_*d*_) of the Ca^2+^ indicator [46,56]. An indicator senses calcium by binding free calcium. In the wider context of buffering effects, calcium binding leads to the removal of free calcium, just as a pump would. The ratiometric cameleon sensor YC2.1 reports a faster Ca^2+^ spike decline and has a larger K_*d*_ for Ca^2+^, than the dye Oregon Green used with the reference Texas Red. Using realistic K_*d*_ appropriate to each sensor, Granqvist et al [56] were able to fit spikes detected with the different sensors. Besides the different K_*d*_, Texas Red is a reference dye insensitive to calcium, while CFP intensity decreases with calcium binding. More detailed spike shape analysis could consider the effects of comparing a ratiometric with a pseudo-ratiometric technique, in particular in very noisy images.

Using similar calcium sensors, spike shape analysis can be applied to detect Ca^2+^ spikes [34] or as yet another characteristic to evaluate Ca^2+^ signatures [25,47,49].

## 4. Imaging Ca^2+^ spiking

The scenario that emerged from imaging studies is of two independent symbiotic calcium signals each predominantly associated with different locations: a Ca^2+^ flux that begins at the root hair tip, and nuclear Ca^2+^ oscillations that initiate from the nucleus periphery. To compare signal intensity in cellular regions with different proportions of calcium sensors, researchers use ratiometric sensors.

In live imaging, the fluorescence signals are often very weak, but options to amplify the signal and obtain an image with a good signal-to-noise ratio (SNR) may conflict with requirements to keep the specimen healthy and optimise the spatiotemporal resolution. Several studies [25,34,35,38] adopted confocal microscopy [58], a technique that enhances optical resolution by the use of a spatial pinhole to remove out-of-focus light. In case the fluorescent signal is very weak, it is possible to capture more signal by increasing the size of the pinhole, but the added signal will arrive from out-of-focus planes, blurring the image. In confocal laser scanning microscopy (CLSM) the image is built by a point-by-point scan, and the first and last scanned pixel in an image frame may be separated by seconds or miliseconds. To amplify the signal we can scan a line several times and then average the image into one, but that is done at the expense of the temporal resolution, which in turn can be increased if we decrease the pixel dwell time, but this again will decrease the signal. And if the laser power is increased to amplify the signal, there is a risk of photobleaching and cell damage. To summarise, live imaging is technically difficult and each particular experiment requires different trade-offs. It may not compensate to increase the signal intensity beyond a certain point, and in that case, imaging processing tools need to tackle the effects of a low SNR.

In what follows, we will show how imaging processing options impact the visual appearance and interpretation of the imaging evidence. We will use examples of calcium signals induced by 10^−9^ M Nod factor in *Medicago truncatula* root hairs. We imaged the root hairs using a confocal microscope (Zeiss Axio Imager Z2 upright microscope, LSM 780) and, as Ca^2+^ sensors, we used members of the yellow cameleon (YC) family of genetically-encoded calcium sensors, specifically YC2.1 (Fig. 4 and 5) and YC3.6 (Fig. 6 and 8). The YC2.1 and YC3.6 cameleons are constructed with a fixed 1:1 stoichiometry of a single donor and acceptor, CFP and YFP, attached to a calcium binding molecule (see Fig 2). As their relative abundance is the same, the ratio between YFP and CFP should in theory be independent of the expression level, with an increase in the ratio of emission at the two wavelengths (YFP/CFP) indicating a relative increase of the Ca^2+^ concentration.

**Figure 2.**
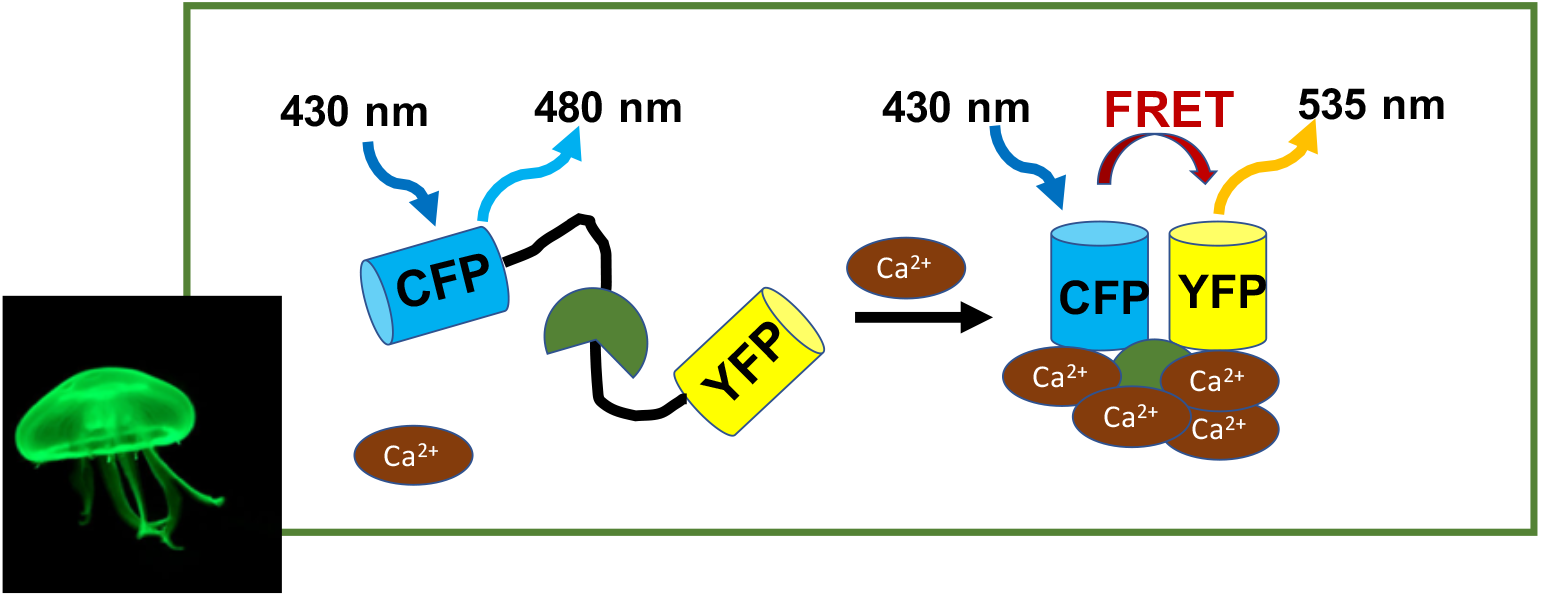
The calcium sensors yellow cameleons shift their emission wavelength upon Ca^2+^ binding. The jellyfish *Aequorea victoria* is the source of the green fluorescent protein (GFP). Yellow and Cyan fluorescent proteins (YFP and CFP) are mutated forms of GFP that absorb and emit light at matching wavelengths, such that the excitation wavelength of YFP is the emission wavelength of CFP. Yellow cameleons are constructed by attaching YFP and CFP to a Ca^2+^-binding domain (green object represents the pair CaM/M13 [24]). When calcium concentration increases, they are brought closer together and experience fluorescence resonance energy transfer (FRET). At the level of a single yellow cameleon construct, a rise in Ca^2+^ concentration is indicated by a drop in the cyan fluorescent protein (CFP) intensity at the same time that the yellow fluorescent protein (YFP) intensity increases.

However, digital images are made up of pixels, and the intensity of a pixel is composed of the signal from the sensor plus a random number corresponding to detector or sample noise. A general rule is that the proportion of signal-to-noise ratio (SNR) is smaller for weaker signals [26]. The challenge is that the cellular regions to compare (nucleus vs tip, or nuclear periphery vs interior) may have very different signal-to-noise ratios. If this is not dealt with, results may be misleading.

Photobleaching is a common feature in fluorescence microscopy, weakening the signal over time as the fluorophores start losing the ability to fluoresce. But in live imaging, the ever-changing SNR appears more complex than a monotonous decay and can pose an hidden complication. Figure 3 shows an idealised representation of any generic cellular object and its corresponding microscopic digital image. The signal — and thus the SNR — depends on continuous cellular motions and shape changes. If our region of interest (ROI) is static while the cellular object moves, the signal/SNR fluctuates alongside variations of the sensor-labeled cytoplasm/nucleus in view. Disregarding shape effects, the problem could be solved by creating a binary signal/noise image, and restricting measures to the dynamic ROI classified as signal. However, we need to take into account the shape of the object, and in particular its edges and interfaces. Any cellular object (eg, nucleus or cytoplasmic strands) is delimited by relatively smooth edges, and the corresponding signal intensity in its 2D representation will be weaker near edges: see the 2D signal corresponding to the sharpest edge at the left (panel A, Fig. 3). Furthermore, often movements and shape changes go together, and all this can be made even more complicated by an uneven local noise from autofluorescent elements. In the simplest case, edges and dynamic cellular regions have a lower SNR.

**Figure 3.**
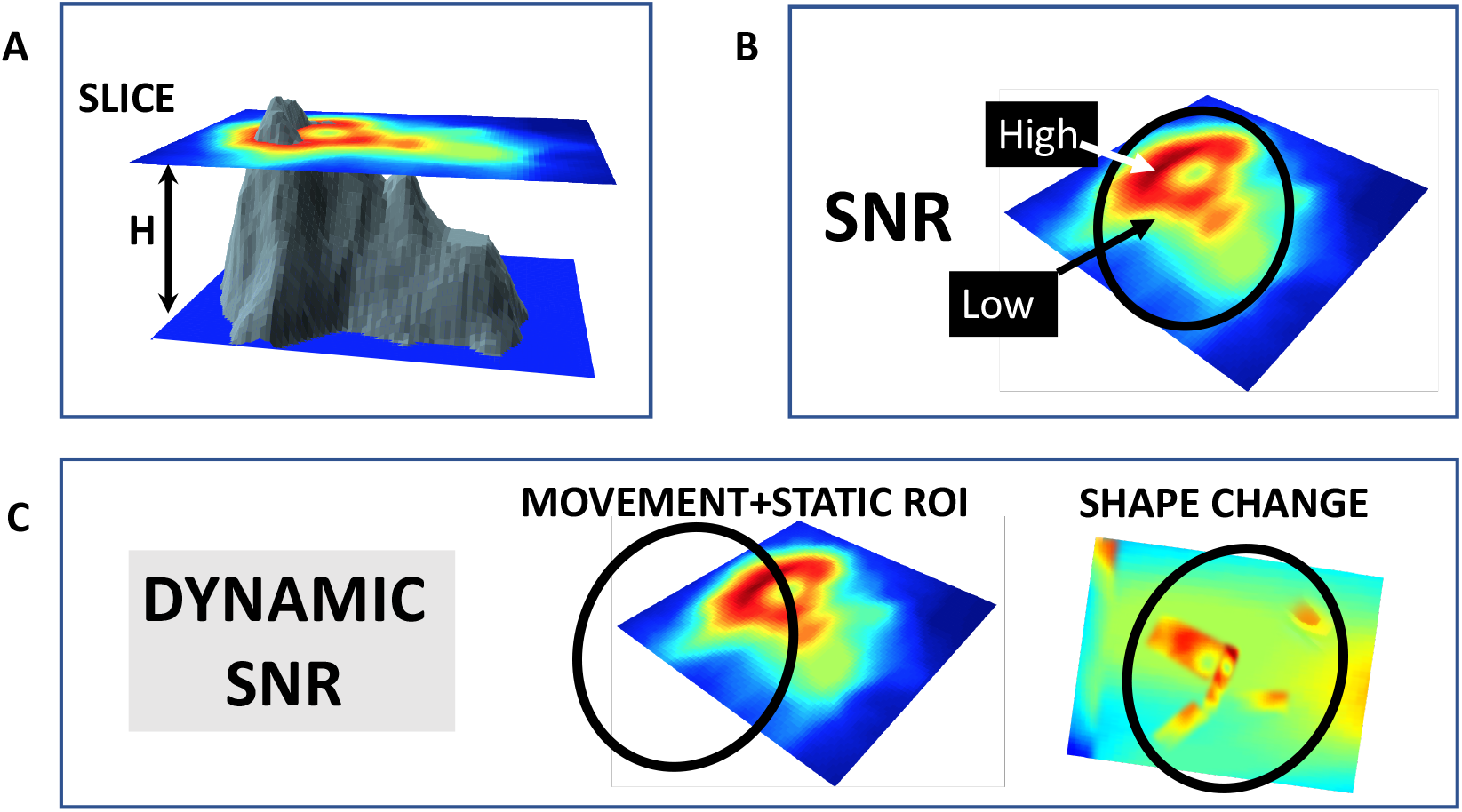
The dynamic nature of the signal-to-noise ratio (SNR) in live imaging. Panel (**A**) In confocal microscopy, the focal plane is defined by the top of the position of the confocal slice through the object (grey). This slice has a small thickness H, which determines how much out-of-focus light is detected. Panel (**B**) The SNR is higher where the signal is stronger: if the sensor is evenly distributed, the SNR at a given point reflects the depth/shape of the object at that point. Panel (**C**) In living samples, the SNR is constantly changing: it decreases if the signal is lowered because the object moves out of a statically defined region-of-interest (ROI, oval), or if the shape of the object changes in certain ways, eg becoming less compact and thus increasing weak edge effects. The idealised cellular object was created from the R volcano dataset [59], using the plot3D R package [60].

With this in mind, we will revisit two phenomena: (1) Ca^2+^ release from the nuclear periphery (Subsection 4.1), and (2) relation between intranuclear and cytosolic Ca^2+^ spiking (Subsection 4.2). Then we will see what imaging can tell us about the relation between perinuclear Ca^2+^ spiking frequency and other signals at the tip (Subsection 4.3). The images we use are not meant to be representative of all symbiotic Ca^2+^ spiking. Instead, they serve to highlight potential pitfalls in the interpretation of images, and to suggest questions for further investigation.

### 4.1. Do we see intranuclear calcium being released from the nuclear periphery?

From other lines of evidence besides imaging, it appears clear that Ca^2+^ is released at the nuclear periphery [35,51,52]. Nevertheless, imaging remains important to understand the scales of the Ca^2+^ spatiotemporal signatures and to pin down the mechanisms of interaction between all the elements involved in Ca^2+^ signalling.

Sieberer et al [34] suggested that the rapid increase in Ca^2+^ concentration observed at the start of a Ca^2+^ spike happens near the nuclear envelope, while the results of Capoen et al [35] show that the nuclear periphery remains brighter throughout the entire spike, as if Ca^2+^ never totally engulfed the entire nucleus. Curiously, the nucleus periphery tends to be brighter even during periods of rest. In Sieberer et al [34] even in between spikes we can observe calcium hotspots along the nuclear border or, more rarely, inside the nucleus. Moreover, in Capoen et al [35], even when the nucleus is not exposed to NF, the border region appears brighter than the interior, although not as bright than when NF is present and Ca^2+^ is spiking. Why, in both studies [34,35], does the nuclear periphery appear brighter even when a Ca^2+^ spike is not imminent? A hypothesis could be that calcium levels were building up at the nuclear border until a threshold is reached when calcium bursts into the nucleus, as long as enough resting time has passed and the right stimulus (NF) is present. However, we cannot confirm or discard this hypothesis before addressing the fundamental problem of how to reliably compare different parts of one image that have different SNRs. To image Ca^2+^ diffusion is not merely a matter of spatiotemporal resolution: as a pre-condition, the Ca^2+^ signal needs to be strong enough in different nuclear regions.

We will look closer at this problem using an example. Figures 4 and 5 refer to the same *M. truncatula* root hair exposed to Nod factor, imaged with a confocal microscope. The calcium indicator is the nuclear-targeted yellow cameleon YC2.1. The image I_1_ of Figure 4 shows the YFP and CFP intensity distributions at the peak of a Ca^2+^ spike. Both YFP and CFP signals are weaker at the nucleolus region and also near the nuclear border. The signal intensity is better captured by a microscope at the centre of the focal plane, weakening the signal at the nuclear edges if the nucleus is at the centre of the image. But even if the nucleus is not at the centre of the image, on a 2D confocal slice the nucleus periphery will show a weaker signal simply because of edge effects (Section 4, Fig. 3). At this point we have a choice before calculating the ratio YFP/CFP.

**Figure 4.**
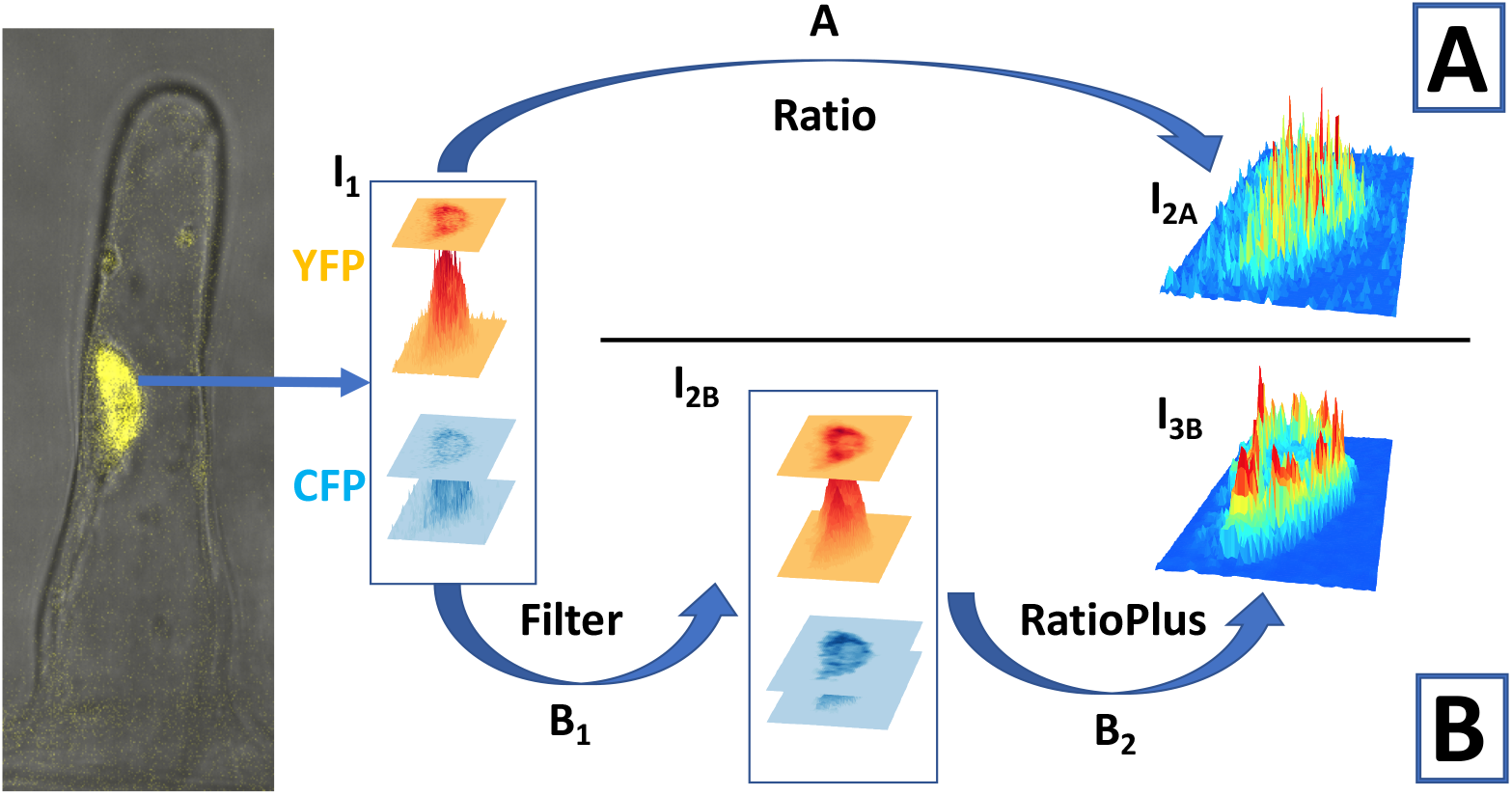
Choices in image processing alter the final ratio image, especially at its edge. The position of a nucleus (yellow) in a *M. truncatula* root hair is shown in the bright field image of the root hair (grey), merged with the YFP image. The path A shows what happens when images are not processed, while the path B attempts to follow the image processing steps reported in Sieberer et al [34]. The 2D microscope image of the YFP and CFP intensities at the nucleus reflects the 3D structure of the nucleus, I_1_ and I_2*B*_. The ratio YFP/CFP images are I_2*A*_ and I_3*B*_. We use the nuclear localised Ca^2+^ sensor YC2.1.

**Option A**, upper panel of Fig. 4 illustrates the ratio image I_2*A*_ obtained without any previous imaging processing. It appears that the Ca^2+^ concentration is largest at very small scattered hotspots, some closer to the nuclear edges and some inside (near the nucleolus). However, this picture of Ca^2+^ distribution is not reliable because noise from fluorescent images is a much larger component of small values [26,27], and the ratios can become artificially large simply because of the small number of photons involved with high uncertainty. Among the different types of noise in fluorescence microscopy [26,27], shot noise, which originates from the particle nature of light, is the fundamental limit of the signal-to-noise ratio [27]. Shot noise follows a Poisson distribution, for which the standard deviation is equal to the square root of the expected value: although its absolute value decreases with the signal intensity, the SNR decreases even faster. If we measure a value within one standard deviation of the true Ca^2+^ signal, the ratio between equally large true signals of, for instance, 400 photons will be measured within the range [0.9 − 1.1], while between true weak signals of 4 photons will be measured within [0.3 − 3], from one third smaller to up to almost 3 times larger. Visually, this could appear as fluctuating hotspots at the nuclear periphery/nucleolus, even in between calcium spikes, and even if we assume for simplicity that the YFP and CFP intensities are the same in those periods of rest (one of the FPs is usually less bright). Therefore, some adjustment should be applied before doing ratios.

**Option B**, lower panel of Fig. 4, attempts to follow the recipe provided by Sieberer et al [34], as closely as we can. The first step is to apply a median filter (step B_1_ in Fig. 4). The median filter removes isolated pixels, either outlier noise or small signal details. Compared with Gaussian blurs or mean filters, it is effective at preserving edges when the original image shows a sharp decline in intensity at the border of the object. We can observe its effects if we compare (Fig. 4) the YFP and CFP signals, before (I_1_) and after (I_2*B*_) filtering. Around the edges, the probability of the signals being outliers is greater, and filtering will not raise intensities at every pixel all around the nuclear periphery. This becomes a problematic region where artefacts arising from the ratio between small random numbers can occur. The nuclear boundary risks being implicitly determined by the statistics of the background noise, which may even vary at different sides of the nucleus (compare YFP, CFP and YFP/CFP nucleus shape, Fig. 5). For the final step before computing the ratio (step B_1_ in Fig. 4), we followed the formula provided by the RatioPlus ImageJ plugin [61,62]. This plugin calculates the pixel-by-pixel ratio after background subtraction and clipping (if after background subtraction the pixel intensity is lower than the clipping value for that image, it is set to zero before ratio calculation) [61]. The image frame is assumed to have a homogeneous background, whose value we chose using the intensities at regions away from the nucleus. We also assume that the background does not change with time, as the plugin can be used on a stack of images simultaneously. To select the clipping value, we take the difference between the measured minimal intensity and the background intensity and divide it by two. Unsurprisingly in our example, after the filtering and RatioPlus steps the final ratio image (I_3_*B*, Fig. 4) shows a much clearer brighter border, compared to the ratio image I_2*A*_

**Figure 5.**
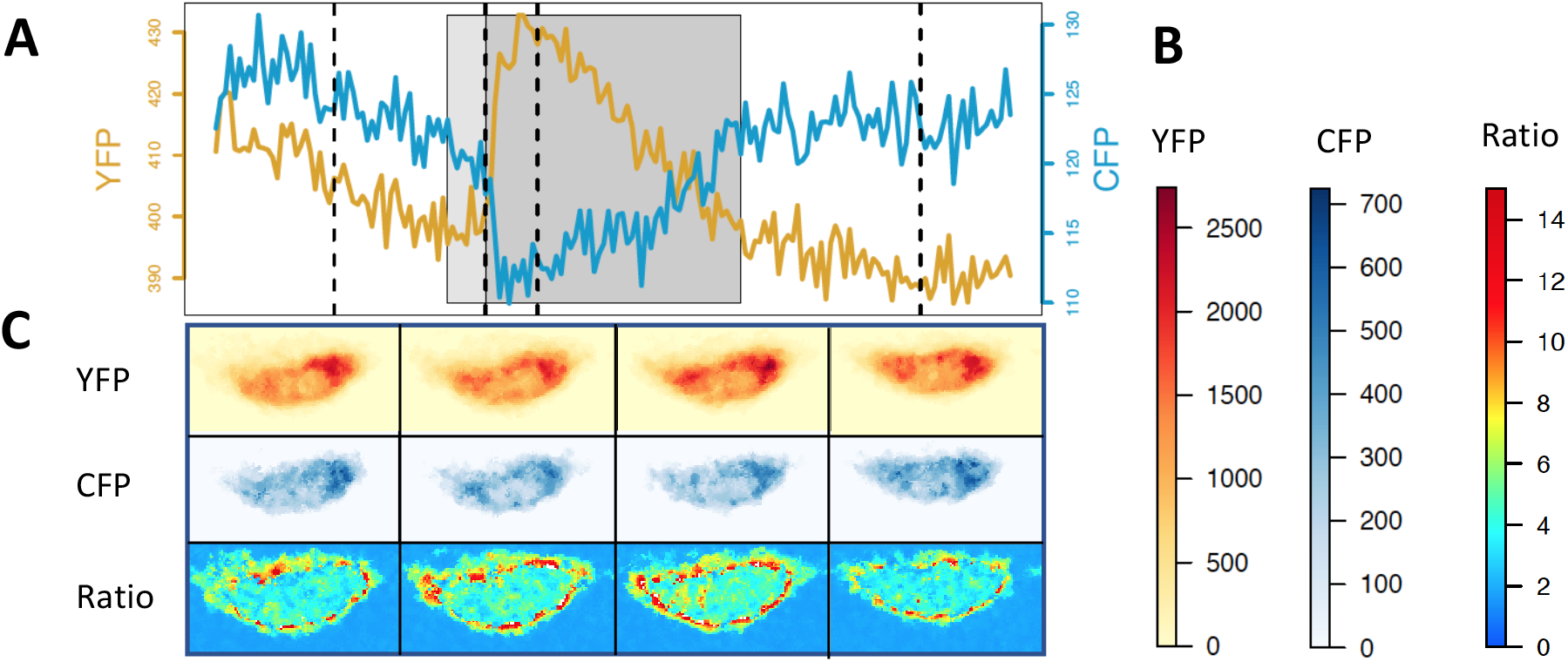
Signal enhancement at the nuclear periphery during a Ca^2+^ spike, when adopting image processing option B of Fig. 4. Panel (**A**) During a Ca^2+^ spike (grey rectangle), the CFP intensity (blue) decreases while YFP intensity (red) increases. Panel (**B**): Colour-bars show that the cyan fluorescent protein (CFP) is less bright than the yellow YFP. Panel (**C**): The signal from YFP and CFP is not uniformly distributed across the nucleus and it is weaker at the periphery. These intensities are presented after filtering, background subtraction and clipping (path B in Fig. 4). Ca^2+^ spike data recorded every 0.37 seconds, and each of the four sequential images for YFP, CFP and Ratio in Panel (**C**) correspond to the vertical dotted lines in Panel (**A**). The Ca^2+^ sensor was the nuclear localised YC2.1.

Such edge artefacts will further be exaggerated if CFP is less intense than YFP, and at the time of a Ca^2+^ spike. The periphery of the nucleus reported by the CFP signal will appear disproportionally shrunk by the filter+background subtraction procedure (see colour-bar scale in Fig. 5). Then, during a Ca^2+^ spike, the CFP signal intensity drops, further worsening its SNR. That can artificially enhance the ratio YFP/CFP near the periphery of the nucleus during a Ca^2+^ spike (Fig. 5).

The chosen [34] ImageJ RatioPlus plugin assumes a very accurate definition of the region of interest [63, 64], such that neither CFP nor YFP signals fall below the intensity threshold that defines background noise. Starting from the same image, in Figure 4, we see two different spatial distribution of Ca^2+^ signals, I_2*A*_ and I_3*B*_. Neither result is the true one, as neither of them solves the problem of compensating for different spatial distributions of the signal-to-noise ratio, but Image I_3*C*_ also shows additional edge enhancements.

We do not know how important were the edge effects discussed above for the images presented in the literature [34,35]. However, unless they are addressed with appropriate statistical tools, the Ca^2+^ levels measured near the periphery of the nucleus will always be biased, to a degree that will depend on the depth of field in the confocal microscope, the fluorescence levels of the specimen, the background values, etc. To bypass the problem of comparing regions with such different SNRs, we could measure signals well inside the nucleus (bar the nucleolus region) and infer the direction of calcium propagation from diffusion profiles. That would possibly imply imaging at even finer levels of spatiotemporal resolution than used so far [34,35], as a clear diffusion profile becomes unclear once calcium is inside the nucleus.

### 4.2. Can we see how apical and perinuclear Ca^2+^ oscillations are related?

Symbiotic Ca^2+^ spiking is nowadays firmly associated with the nuclear and perinuclear region. In imaging studies, spikes appear delayed with distance from the nucleus [31,38]. Often, the shape of cytosolic calcium spikes degrades and dissipates farther from the nucleus. This makes cytosolic Ca^2+^ spiking appear almost like a casual byproduct, and if it encoded a message, it would be discarded as meaningless. However, how does a worse SNR at the root hair tip alter the appearance of Ca^2+^ spikes at the tip?

Unless the root hair is so young that it still exhibits a cytoplasmic plug at the tip, the signature of Ca^2+^ spiking far from the nucleus is complicated by the impact of cytoplasmic streaming, which constantly rearranges the distribution of Ca^2+^ sensors seen in a static region-of-interest. A clearer signal is expected where Ca^2+^ sensors continuously occupy a relatively large and stable region like the nucleus and the cytosol surrounding it. Can the apparent degradation of Ca^2+^ signals near the root hair tip be an artefact of cytoplasmic streaming? While Ca^2+^ release is indicated by an increase in the ratio YFP/CFP, intracellular motions can possibly cause changes in the SNR that affect that ratio. When motions within the cell are significant and happen at similar time scales as Ca^2+^ release, it may be challenging to disentangle the causes of small YFP/CFP changes. We illustrate this challenge with an example of a *M. truncatula* root hair exposed to NF, now using the cytosolic/nuclear Ca^2+^ sensor YC3.6 (Fig. 6).

**Figure 6.**
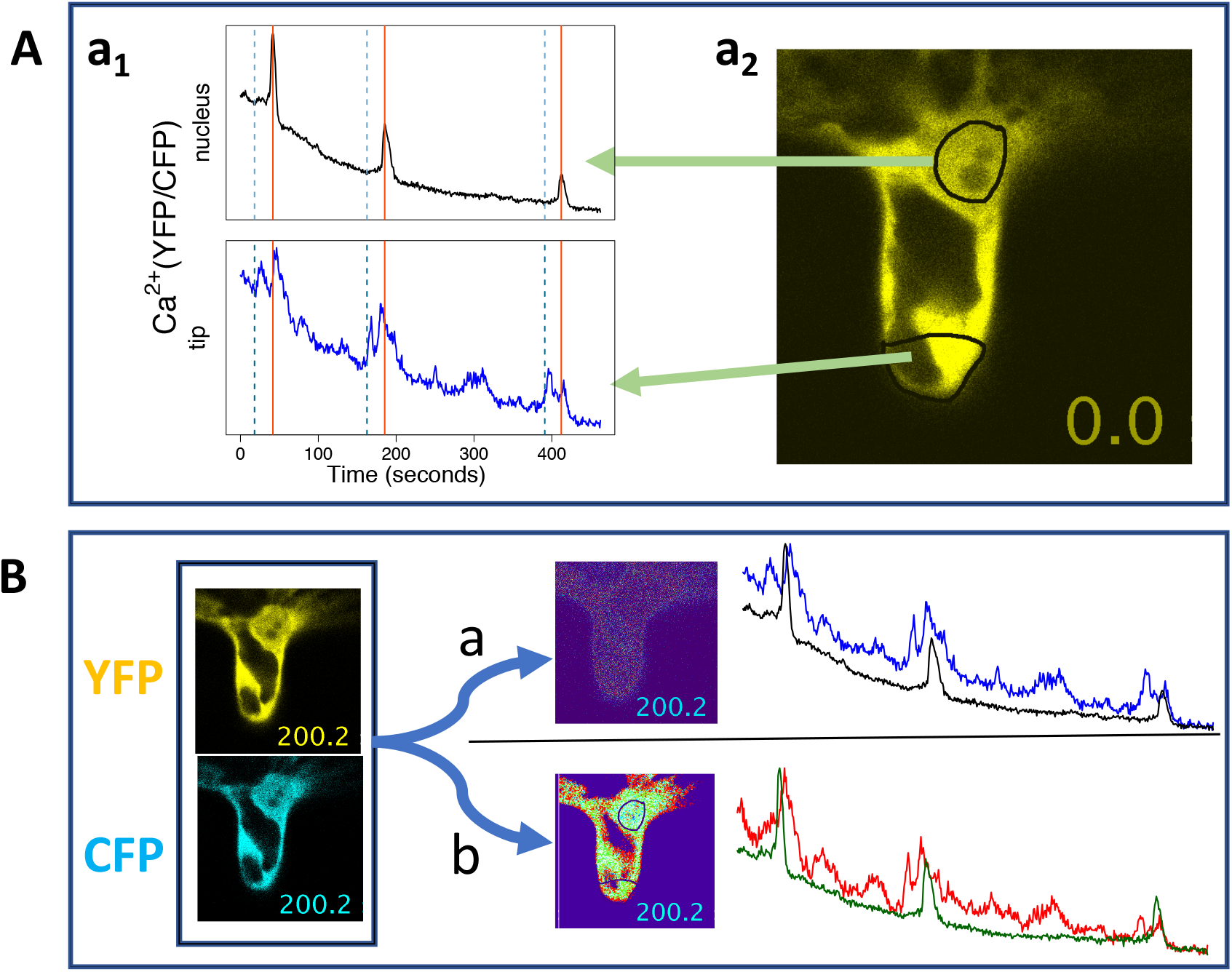
Apparent apical and nuclear Ca^2+^ spikes are related in a complicated way. Panel (**A**), *a*_1_, shows oscillations in perinuclear and apical YFP/CFP ratios, corresponding to the statically-defined ROIs observed in the image of the root hair YFP signal (*a*_2_), where the black region corresponds to the sensor-free vacuole. Panel (**B**) shows the impact of imaging processing on ratio images (*t* = 200.2 sec) and on the YFP/CFP time series. Option *a*, panel (**B**), corresponds to the use of RatioPlus [61] plugin without background subtraction and without previous filtering of the YFP and CFP images. Option *b*, panel (**B**), corresponds to the application of a 2 pixel median filter on the YFP and CFP images, followed by the use of RatioPlus with background subtraction and clipping. Data recorded every 0.79 seconds, and the Ca^2+^ sensor was the cytosolic/nuclear localised YC3.6. Note that although the times at the first and last pixels in an image frame are not exactly the same (because of line scanning), the relation between nuclear and apical traces is not significantly affected by the scanning starting position: if we translate the tip traces by one time point, we don’t see any significant difference.

We present the Ca^2+^ (YFP/CFP) traces, panel A *a*_1_, Fig. 6, without any conventional pre-processing: no denoising and no background and trend removal [5,8,65]. As is typical, the Ca^2+^ spikes are much sharper at the perinuclear region. However, there are a few atypical observations. The larger peaks of the apical Ca^2+^ spikes are not always delayed relative to the corresponding perinuclear Ca^2+^ spikes. Furthermore, each Ca^2+^ spike at the tip seems composed of a previous spike that happens before the major perinuclear Ca^2+^ peak (vertical dotted lines, panel A *a*_1_, Fig. 6). In fact, the *precursor spike* coincides with a gradual growth of perinuclear Ca^2+^ concentration, before the characteristic steep spike rising phase that is reported in the literature.

Note that Ca^2+^ signals were imaged in this root hair more than half an hour after exposure to NF, so even if there had been a previous apical Ca^2+^ flux (not observed), at this point we wouldn’t expect to observe any interference with the typical Ca^2+^ spiking associated with the nucleus. On this account, is the coincidence between the gradual start of the perinuclear Ca^2+^ spikes and the existence of *precursor* apical Ca^2+^ spikes an artefact? In other words, do the apical YFP/CFP ratio changes indicate real Ca^2+^ transients?

If some features of the apical spikes are artefacts caused by the changing distribution of the Ca^2+^ sensors, then imaging processing steps that alter the spatial distribution of the YFP and CFP signals could potentially alter the appearance of the time series. Panel B, Fig. 6 illustrates the effect of filtering and background subtraction (option *b*), as compared to simple pixel-by-pixel ratio (YFP/CFP) calculation (option *a*). The first observation is that without filtering the ratio image appears so noisy that we can hardly recognise the cytosol/vacuole interface. Secondly, we see that filtering and background subtraction changes the apical YFP/CFP traces to reveal *bumps* with size and time scales comparable to the apparent apical Ca^2+^ spikes. Given this, it would be unclear how to adapt the usual pre-processing steps, such as trend removal, to these time series. In fact, with this option *b*, the existence of any apical Ca^2+^ spikes looks uncertain. However, it is still possible to distinguish apparent apical double Ca^2+^ spikes in both (*a* and *b*) cases, which coincide with the perinuclear Ca^2+^ spikes.

To investigate further, we focus on the second spike before t=200 s, Fig. 7, panel A. Whereas the nuclear ROI seems to capture a fairly stable large area occupied by the sensor (with the exception of the nucleolus), the ROI at the tip largely coincides with the vacuolar region. Zooming on the tip and the nuclear region, panel B, Fig. 7, we observe that when the *precursor* apical peak (point 2, t=168.0 sec) occurs, the brightest (red) pixels in the ratio image are seen in places where there are almost no YFP and CFP signals. This suggests that the apical and perinuclear YFP/CFP peaks may be caused by different reasons: while perinuclear high ratios may reflect a true increase in Ca^2+^ concentration, apical high ratios may have been artefacts of the division between small random numbers, similarly to what we discussed previously (Subsection 4.1). In fact, the composite apical Ca^2+^ spike coincides with a generic drop of both the YFP and CFP intensity, although YFP grows at the time of the major spike (panel C, Fig. 7). Note also that a gradual Ca^2+^ growth before the perinuclear Ca^2+^ spike coincides with a decay of both YFP and CFP and also with the start of the major apical Ca^2+^ spike (grey polygon, panel C, Fig. 7).

**Figure 7.**
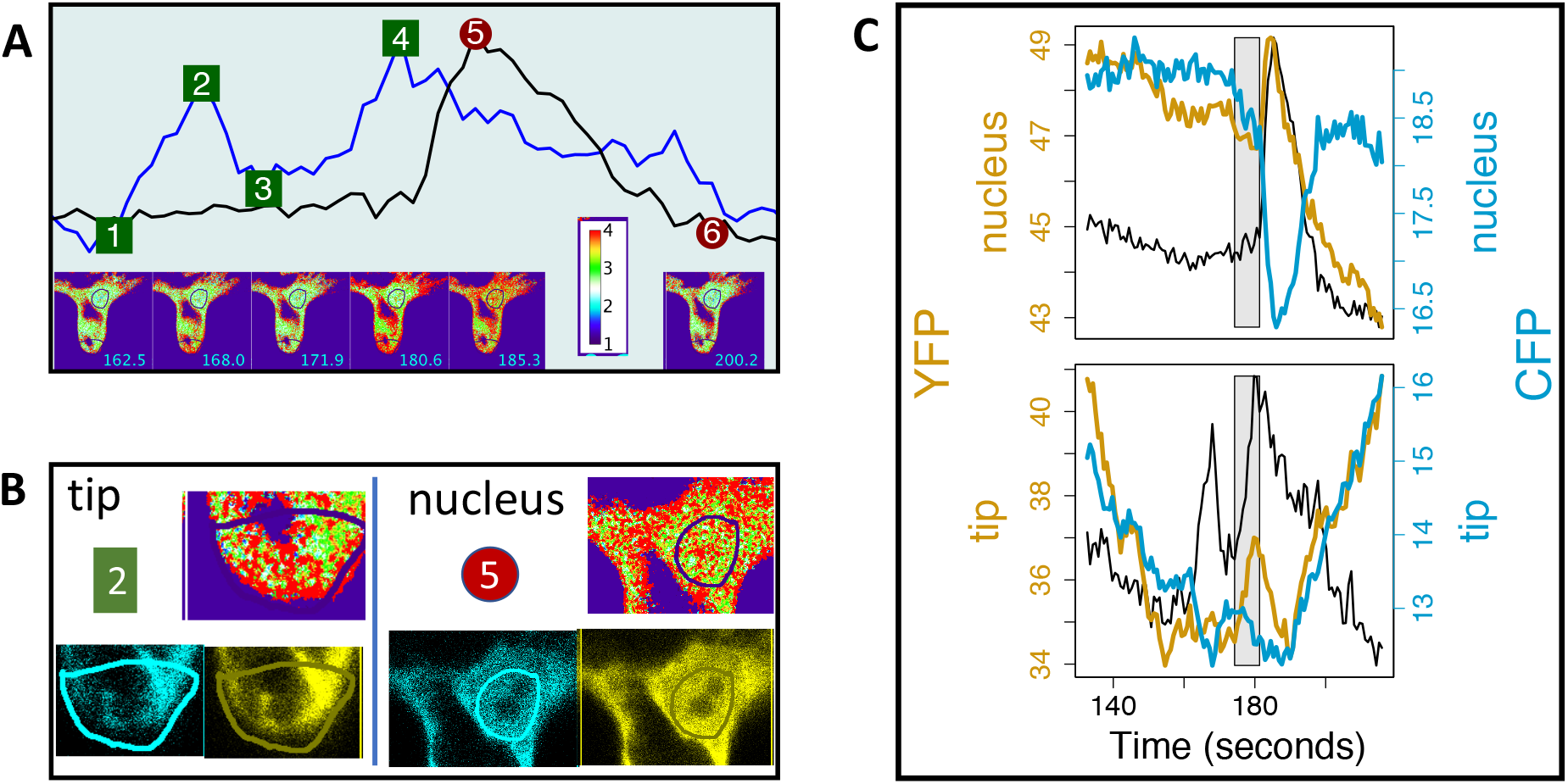
Increases in the ratio YFP/CFP may be caused by noise, more so in the apparent *precursor* apical Ca^2+^ spike. Panel (**A**), shows the *second* (before t=200 sec) perinuclear Ca^2+^ spike (black), superimposed on the corresponding apparent double apical Ca^2+^ spike (blue). Panel (**B**) shows a snapshot of the YFP, CFP and ratio images at the time of the first apical peak (t=168.0 sec, 2) and of the perinuclear spike (t=185.3 sec, 4): large intensity (red) in the apical ratio image happen at pixels where the apical YFP, CFP images hardly show any signal. Panel (**C**) shows coincidences between changes in parallel and antiparallel CFP and YFP intensities (grey polygon). Data recorded every 0.79 seconds, and the Ca^2+^ sensor was the cytosolic/nuclear localised YC3.6.

It is clear from this example that changes in YFP and CFP intensity can be related in a complex way to Ca^2+^ spiking. The existence of perinuclear Ca^2+^ spikes is clearly not an artefact, as it coincides with antiparallel YFP and CFP changes, and the same can be said for the second peak of the apical Ca^2+^ spike, panel C, Fig. 7. But doubts remain about the apparent gradual start of the perinuclear Ca^2+^ spike, and the *precursor* apical YFP/CFP peak. We can indirectly infer movements and shape changes from parallel (instead of antiparallel) changes in YFP and CFP intensities, panel C Fig. 7. It could have happened that cytoplasmic streaming led to a decrease of the signal intensity, and a few CFP pixels have fallen below the detection threshold. The partial loss of CFP signal would then artificially increase the YFP/CFP ratio to reveal apparent double apical Ca^2+^ spikes. As cytoplasmic streaming also affects the shape and motions of the nucleus, the apparent gradual start of the perinuclear Ca^2+^ spike could also be an artefact.

However, even if some YFP/CFP ratio peaks don’t indicate true Ca^2+^ changes, it is intriguing to observe them so close to each perinuclear Ca^2+^ spike. Can cytoplasmic streaming be biologically related to the timing of Ca^2+^ release? While previously we saw how results obtained with the use of ratiometric sensors demand a careful interpretation, next we will see how ratiometric sensors can provide additional interesting information, besides the information about Ca^2+^ transients.

### 4.3. Ca^2+^ signals and other signals at the tip

Already several components of the Ca^2+^ nuclear signalling machinery have been identified, which according to mathematical models [10] can be enough to sustain periodic oscillations, once Ca^2+^ spiking is initiated. These mechanistic models are valid if several unknown parameters are tuned adequately. Can imaging reveal any other signal involved in sustaining or initiating Ca^2+^ spiking?

Figure 8, panel A, shows perinuclear regular Ca^2+^ oscillations, imaged using the nuclear/cytosolic sensor YC3.6, in *M. truncatula* root hair. At the tip, spiking is hardly visible, and we can barely distinguish a few coincidences between perinuclear Ca^2+^ spikes and small apical Ca^2+^ increases. Strikingly, a much more clear relation is found between perinuclear Ca^2+^ spiking and parallel changes in intensity of YFP and CFP at the tip: they appear synchronised, panel C, Fig. 8. Naturally, we don’t observe oscillations of the YFP and CFP intensities around a certain constant value, as that would have no biological justification. But we do see a clear coincidence between perinuclear Ca^2+^ spikes and the speed and/or direction of change in apical YFP and CFP intensities. Since the regions-of-interest (ROI) were statically defined at start, these changes are likely to be related to cytoplasmic streaming at the tip, panels A and C Fig. 8.

**Figure 8.**
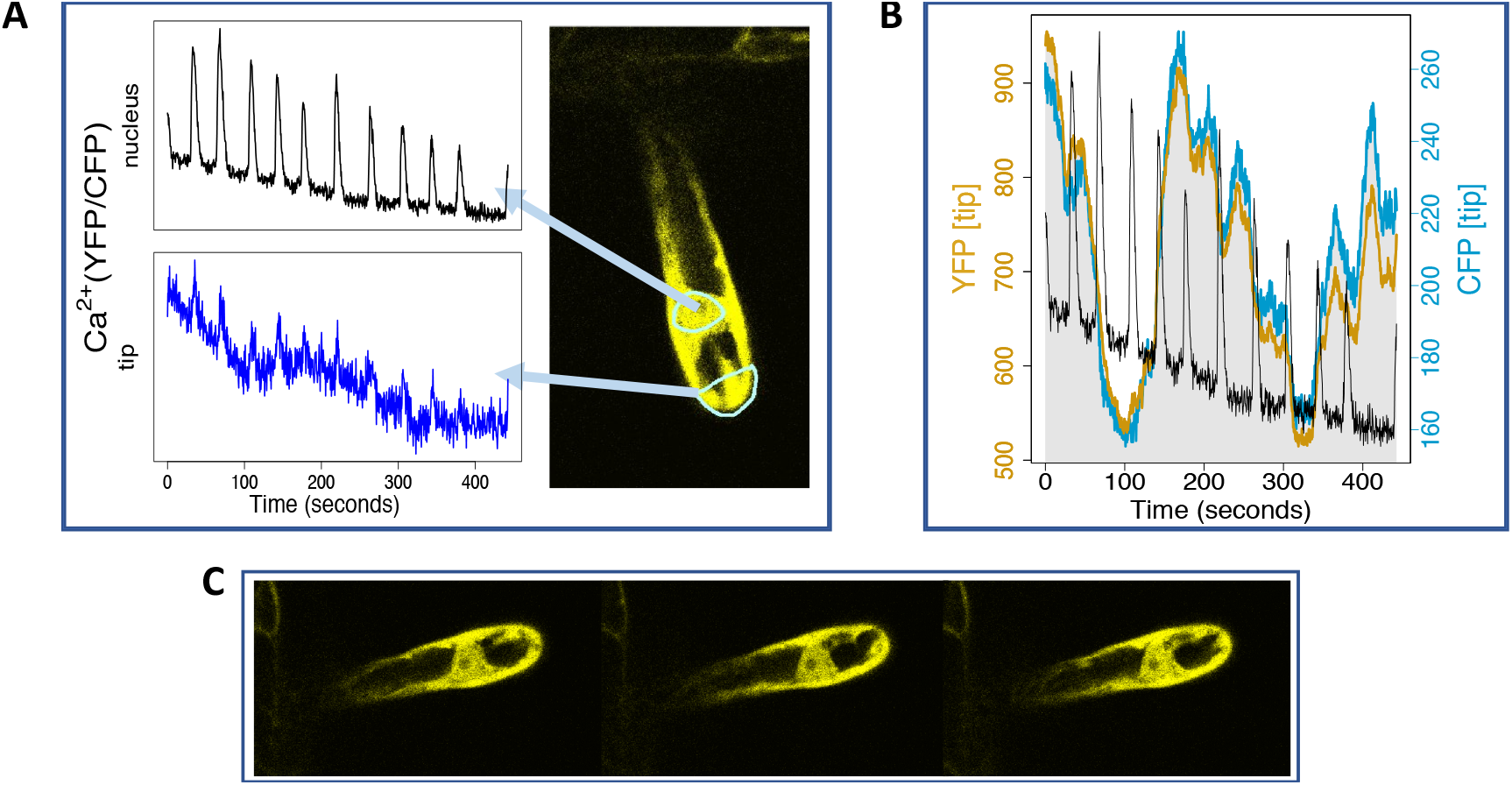
Is the Ca^2+^ signature related to cytoplasmic streaming? Panel (**A**) shows perinuclear and apical Ca^2+^ spikes, corresponding to the statically-defined regions shown in the YFP signal. Panel (**B**) shows that perinuclear Ca^2+^ spiking occurs when there is a parallel change in the YFP and CFP intensities at the tip region. Examples of how the appearance of cytoplasm at the tip changes is seen by comparing Panels (**A**) and (**C**), which show the YFP images at different times. For the time series in panels (**A**) and (**B**), time points were taken every 0.44 seconds. The Ca^2+^ sensor was the cytosolic/nuclear localised YC3.6.

This example reinforces the idea, advanced in Subsection 4.2, that there is a relation between motions at the tip region, and nuclear Ca^2+^ spiking. It is presented to stimulate questions. The nucleus and the tip are connected by cytoplasm or cytosolic strands with decreasing degrees of density as the root hair grows older. Although symbiotic Ca^2+^ spiking is induced by NF, and root hair cells can exhibit cytoplasmic streaming without any relation to NF, Sieberer et al [66] found that patterns of cytoplasmic streaming change with exposure to NF. Can Ca^2+^ signatures, which are related to the development stage of the root hair [46], be related to specific patterns of cytoplasmic streaming? Is each calcium spike related to mechanical stimuli at the nuclear border? If each spike is related to a rearrangement of the cytoplasm distribution, can this also explain why spikes appear so much noisier at the tip? This could possibly be the case if changes coincided with the temporary appearance of a more web-like cytoplasmic distribution, as this spread would increase the number of noisier cytoplasm/vacuole interfaces at the tip. Answering these questions would require new images to build robust statistics, and also advanced imaging processing methods, which would address the issues raised above.

To begin to disentangle biological from spurious correlations, we could start to systematically investigate whether Ca^2+^ release and cytoplasmic streaming appear related only at the times when the SNR decreases. Another point, is that *cytoplasmic streaming* should be better characterised. It is not ideal to infer movement from parallel changes in YFP and CFP intensity, as they are affected by a variety of factors, such as photobleaching. Movements and shape changes can, in theory, be characterised easily in binary images. However, in practice, low SNRs can lead to the introduction of additional artefacts when binary masks are created. The decision to create masks or use indirect measurements depends on the balance of which artefacts can be better canceled, for each particular case. It is also important to systematically compare results obtained with different sensors (GECI or dyes) and with different imaging techniques (eg, confocal or wide-field microscopy.)

## 5. Conclusion

We have considered some of the fundamental imaging studies that have informed the current understanding of Ca^2+^ signalling during the initiation of legume-bacteria symbiosis. Many of the studies combine imaging with other lines of evidence, from genetic studies to mathematical models. However, each of them contributed to the construction of a coherent narrative of symbiotic calcium signalling based on imaging. They supported the idea of two symbiotic Ca^2+^ signals: a Ca^2+^ flux at the root hair tip and nuclear Ca^2+^ oscillations with different frequencies characteristic of different symbiotic phenomena.

Historically, fluorescence microscopy was a qualitative technique, valuable for stimulation of new avenues of research or for visual illustration of evidence backed by other methods. In recent years, and in parallel with advances in high-resolution microscopy and development of new fluorescent sensors, bioimaging has been fast progressing from direct observation to quantification, with new methods serving to estimate and reduce the uncertainty in the imaging data [16]. Here, we have reviewed the literature, and attempted to identify challenges and potential pitfalls. Regarding the legume-rhizobial symbiosis, one source of uncertainty arises from the comparison between Ca^2+^ signals in cellular regions disproportionally affected by noise: the problematic regions are nuclear periphery affected by low SNR at edge artefacts, and the root hair tip affected by cytoplasmic streaming.

Shedding light into the consequences of image processing approaches, we suggest that Ca^2+^ spiking is not as localised as commonly thought. Regarding intranuclear Ca^2+^, its stronger concentration at the nuclear periphery is not presently confirmed by robust image processing approaches. That does not invalidate the fact that the Ca^2+^ signal machinery is localised at the nuclear membranes; instead, it questions its limited reach. The apparent dissipation of Ca^2+^ spiking with the distance from the nucleus could possibly be explained by an increased SNR at the tip caused by cytoplasmic streaming. A detailed comparison between *degraded* apical Ca^2+^ spikes with *sharp* perinuclear Ca^2+^ spikes reveals unexpected correlations: in some cases, an apparent apical Ca^2+^ spike seems to precede a perinuclear Ca^2+^ spike. However, we note that when measuring small apparent Ca^2+^ transients by ratiometric sensors, sometimes we are unable to rule out artefacts: eg., cytoplasmic streaming can cause the CFP signal to become partially undetectable, leading to artificially high YFP/CFP ratios. Does cytoplasmic streaming merely affect the measurement of Ca^2+^ signals, or does it affect the Ca^2+^ signals themselves, as it sometimes appears to interfere at particular moments, at the start of a spike? We show one example where the most unequivocal relation is not between apical and perinuclear Ca^2+^ signals but between changes in apical cytoplasmic streaming and the frequency of perinuclear Ca^2+^ signals. If confirmed by future studies, this will show that the entire root hair participates in Ca^2+^ spiking throughout its duration.

For the interdisciplinary synergy that is key in bioimaging, biological expertise should be complemented by the skills of physicists, computational scientists and microscopy specialists. Computational methods would help formalise and standardise reproducible imaging protocols, with estimation of uncertainties [67]. Microscopy specialists would help understand the noisy features hidden in the images. The contribution of physicists working in the area of complex systems could help elucidate ideas like cell autonomy: nuclear Ca^2+^ spiking is thought to be cell-autonomous, but per se, frequency mismatch doesn’t imply independence [68], as nonlinear interactions can manifest in unexpected ways. A complex systems approach could also help in determining the precise way (mechanical effects, transport of biochemical stimulus, particular patterns) [69] in which cytoplasmic streaming is — or not — related to the Ca^2+^ spiking rhythm.

### Author Contributions

conceptualization, TVM; methodology, TVM and VNL; investigation, TVM; writing and review and editing, TVM and VNL.

### Funding

This research was funded by BBSRC’s Institute Strategic Programme on Biotic Interactions underpinning Crop Productivity (BB/J004553/1) and Plant Health (BB/P012574/1).

## Acknowledgments

TVM acknowledges Sarah Shailes, Sylvia Singh, Grant Calder and Jongho Sun for advice on plant preparation and confocal microscopy. We acknowledge Richard J Morris and Allan Downie for very helpful comments.

## Abbreviations

The following abbreviations are used in this manuscript:

SNR: Signal-to-noise ratio
*M. truncatula*: *Medicago truncatula*
*S. rostrata*: *Sesbania rostrata*
OG: Oregon Green
TR: Texas Red
Nod: Nodulation
NF: Nod factor
FRET: Fluorescence (or Forster) Resonance Energy Transfer
YC: yellow cameleon
YFP: yellow fluorescent protein
CFP: cyan fluorescent protein
CLSM: confocal laser scanning microscopy

